# A protocol for chemical competence in phytopathogenic *Ralstonia*

**DOI:** 10.1101/2025.10.13.678364

**Authors:** Tabitha C. Cowell, Noah R. Guillome, Matthew L. Cope-Arguello, Neha N. Prasad, Tiffany M. Lowe-Power

## Abstract

Here, we present a protocol for the transformation of *Ralstonia solanacearum* species complex (RSSC) strains by calcium chloride-induced chemical competence. We include a step-by-step protocol with media and solution recipes, quantitative results of the protocol’s efficacy for four of five tested RSSC strains, and an explanation of the logic underlying the protocol development. The protocol involves sequentially treating overnight cultures with a cold calcium chloride solution and a cold calcium chloride/magnesium chloride/glycerol solution, transforming through alternating heat shocks and ice incubations, and recovering cells via an outgrowth in rich medium.

**Data summary:** Supplemental File S1 is the preliminary version of this chemical competence protocol. Supplemental Figures S1 and S2 show transformation efficiency data for pUFR80 knockout plasmids, and heat shock number and temperature variation, respectively.

## Introduction

The *Ralstonia solanacearum* species complex, hereafter “*Ralstonia*”, includes three species of plant pathogenic bacteria, divided into four phylotypes: *R. solanacearum* (phylotype II), *R. pseudosolanacearum* (phylotypes I and III), and *R. syzygii* (phylotype IV) (1). These *Ralstonia* species are in the Burkholderiaceae family and are model organisms for studying bacterial pathogenesis and physiology. Genetic tractability is a key requirement for model organisms. Currently, there are established methods for electroporation, natural transformation, and conjugation for *Ralstonia* (2,3). Here, we developed a protocol for *Ralstonia* transformation using calcium chloride-induced chemical competence. Like natural transformation, chemical competence methods are low-cost and accessible because they do not require specialized equipment and consumables like electroporators or electroporation cuvettes. Here, we present our chemical competence protocol, as well as protocol validation and logic underlying protocol development sections.

## Protocol

### Reagents

#### Media

Media are sterilized by autoclaving. TZC and antibiotics should be added to molten CPG agar cooled to ∼57°C. Common antibiotics and final concentrations for genetically engineering *Ralstonia* are kanamycin (25 μg/mL), gentamicin (10 μg/mL), tetracycline (30 μg/mL), and spectinomycin (40 μg/mL). *Ralstonia* isolates often have natural resistance to the antibiotics chloramphenicol (5), penicillin (5), ampicillin (6), and trimethoprim (unpublished data, Lowe-Power lab).

### Solutions

Solutions should be filter-sterilized using filters with a 0.22 μm pore size and pre-chilled to 4°C prior to use.

### Procedure

After the written protocol, there are notes and recommendations for steps 5, 6, 9, 12, 14, 19, and 20.

#### Culture Growth

1. Streak out *Ralstonia* strain on CPG+TZC agar and incubate at 28°C for 2 days or room temperature (20-22°C) for 3-4 days.
2. Inoculate a single colony into 5 mL CPG broth and incubate overnight in a shaking incubator at 250 rpm and 28°C.

#### Cell Treatment

3. Perform all steps on ice except for centrifugations. Ensure that materials are pre-chilled: solutions and microcentrifuge tubes (1.5 mL and 2.0 mL). Set a water bath to 45°C.
4. Incubate the overnight culture on ice for 10 min (make sure the culture tube is slightly buried in ice so that the full culture volume is chilled).
5. Add 1 mL overnight culture to a chilled 1.5 mL microcentrifuge tube.
6. Centrifuge for 30 s at 13,000 x*g* and pipette off the supernatant.
7. Resuspend cell pellet in 1 mL cold 100 mM CaCl_2_ by pipetting to mix.
8. Incubate on ice for 20 min.
9. Centrifuge for 30 s at 13,000 x*g* and pipette off the supernatant.
10. Resuspend cell pellet in 1 mL cold TG salt solution by pipetting to mix.
11. Incubate on ice for 15 min.
12. Centrifuge for 30 s at 13,000 x*g* and pipette off the supernatant.
13. Resuspend cell pellet in 100 µL cold TG salt solution by pipetting to mix and keep the chemically competent cell suspension on ice until use.

#### Transformation

14. Add plasmid DNA and 100 µL of the chemically competent cell suspension to a chilled 2.0 mL microcentrifuge tube.
15. Incubate on ice for 15 min.
16. Heat shock in a water bath at 45°C for 2 min.
17. Incubate on ice for 15 min.
18. Repeat steps 16-17 twice for a total of three heat shocks and four ice incubations.
19. For the outgrowth, add 500 µL CPG broth and shake the tube horizontally for 4 hours in a shaking incubator at 250 rpm and 28°C.
20. Plate 100 µL of the outgrowth culture onto prewarmed plates containing CPG+TZC media with selective antibiotics. Incubate at 28°C for 2-5 days.

#### Notes and Recommendations

- Steps 5-6: More than 1 mL of the overnight culture can be pelleted if the user requires a greater number of cells, or if the culture is harvested at a low cell density.
- Steps 6, 9, 12: Some cells will be lost after each centrifugation and supernatant removal, but this is not a problem when working with high density overnight cultures (A_600_ > 2). If cells are centrifuged for longer than 30 s, we recommend using a temperature-controlled centrifuge set to 4°C to prevent the cells from warming during centrifugation. We did not yield any transformants after six transformations from a single attempt to scale up “Cell Treatment” (Steps 3-13) by centrifuging in 50 mL conical tubes at 5,000 x*g* for 10 min at 4°C.
- Step 14: When using plasmids that are 8 kb or larger, we recommend using large amounts of plasmid DNA (≥ 1 μg) for higher rates of successful transformation. We chose to move cells to 2.0 mL microcentrifuge tubes in Step 14 because these tubes allow for better circulation of the culture during the outgrowth.
- Step 19: The outgrowth can be carried out for as little as 2 hours. Prolonged outgrowths in excess of 5 hours should be avoided so that each colony on the selection plate is derived from an independent genetic event.
- Step 20: The outgrowth culture can be diluted before plating, and a 1:10 dilution is usually sufficient. The outgrowth culture can also be concentrated by centrifuging and resuspending the cells in a smaller volume prior to plating.

## Protocol Validation

### Bacterial Strains, Plasmids, and Growth Conditions

We developed the chemical competence protocol using the phylotype I *R. pseudosolanacearum* strain GMI1000. The protocol was also tested on additional strains, including the phylotype II *R. solanacearum* strain IBSBF1503, the phylotype IV *R. syzygii* strain PSI07, and the phylotype III *R. pseudosolanacearum* strains UW386 and CMR15 (**Table 4**). All strains were stored as glycerol stocks at −80°C.

We developed this protocol using pBBR1-MCS5 (13) and pSW002 (14) (**Table 5**). pBBR1-MCS5 is a 4.8 kb plasmid that can replicate in *Ralstonia* and confers gentamicin resistance. However, we caution against selecting pBBR1 to genetically engineer *Ralstonia solanacearum* species complex strains. We have anecdotally observed that plasmids with pBBR1 origins of replication (including pUFJ10) sometimes “make the strain sick” yielding colonies with an appearance of reduced exopolysaccharide production and elevated production of diffusible melanin. pSW002 is an 8.2 kb plasmid with a pVS1 origin of replication that can replicate in *Ralstonia* and confers tetracycline resistance. To date, we have not observed aberrant colony morphologies with pVS1-ori plasmids in *Ralstonia solanacearum* species complex strains. Plasmids were stored in *Escherichia coli* as glycerol stocks at −80°C.

Plasmid extractions were performed according to the manufacturer’s instructions with the Zyppy Plasmid Miniprep Kit (Zymo Research, Cat. No. D4037), GeneJET Plasmid Miniprep Kit (Thermo Scientific, Cat. No. K0503), or I-Blue Mini Plasmid Kit (IBI Scientific, Cat. No. IB47172), using the kit’s elution buffer or water to elute the plasmid DNA. Plasmid DNA concentration and purity were assessed using a NanoDrop One (Thermo Scientific).

For growing *Ralstonia*, we used CPG media (4) (**Table 1**). Media were sterilized by autoclaving. Liquid cultures were grown in a shaking incubator at 28°C and 250 rpm. Cultures on agar plates were grown at 28°C. Concentrations and catalogs for antibiotics are listed in “Media”.

**Table 1.**
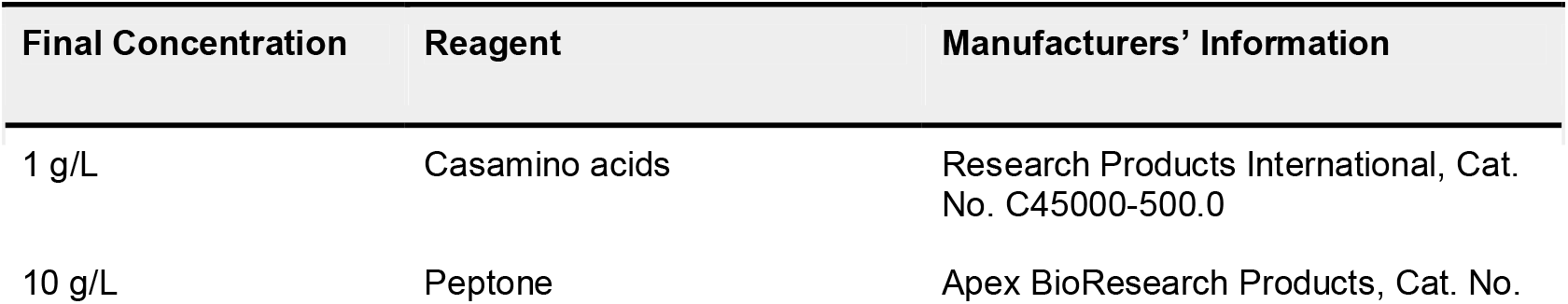
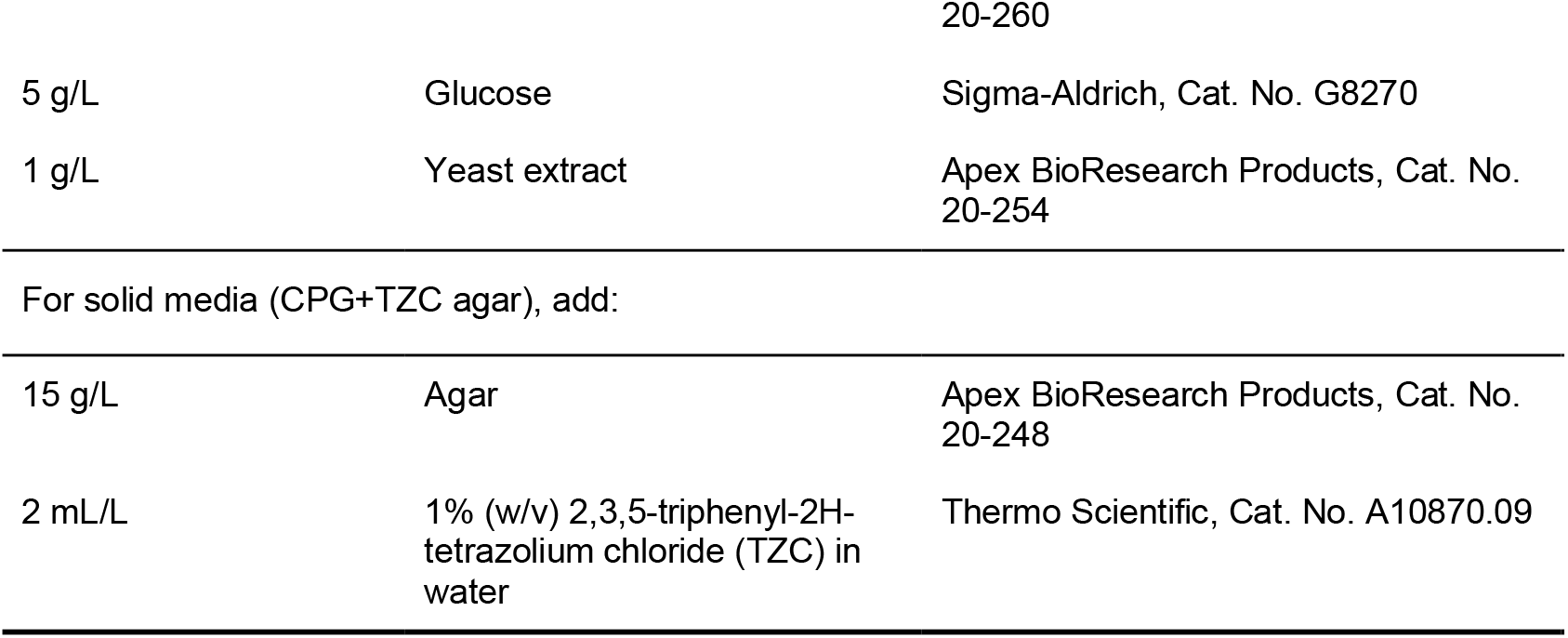
CPG rich medium (4).

There were two media recipes developed for a preliminary chemical competence protocol (Supplemental File S1). The first was a modified Super Optimal Broth (SOB) for *Ralstonia* (ROB), which was composed of 1 g/L casamino acids (Research Products International, Cat. No. C45000-500.0), 15 g/L peptone (Apex BioResearch Products, Cat. No. 20-260), 1 g/L yeast extract (Apex BioResearch Products, Cat. No. 20-254), 12.5 mM KCl (EMD, Cat. No. PX1405-1), 10.0 mM MgCl_2_ (Sigma-Aldrich, Cat. No. M8266), and 10.0 mM MgSO_4_ · 7H_2_O (Fisher Scientific, Cat. No. M80-500). The second was a modified Super Optimal broth with Catabolite Repression (SOC) for *Ralstonia* (ROC), which is ROB with glucose (Sigma-Aldrich, Cat. No. G8270) added to a final concentration of 20 mM (or 0.4% (w/v)). Although glucose induces catabolite repression in *E. coli*, a study recently demonstrated that *Ralstonia* do not have glucose-inducible catabolite repression (15). Both ROB and ROC were filter-sterilized using a 0.22 μm filter before use.

There were two solutions used in the chemical competence protocol. The first was 100 mM CaCl_2_ (**Table 2**). The second was Transformation salts with Glycerol (TG salt solution) (7) (**Table 3**). Both solutions were filter sterilized using a 0.22 μm filter before use.

**Table 2.**
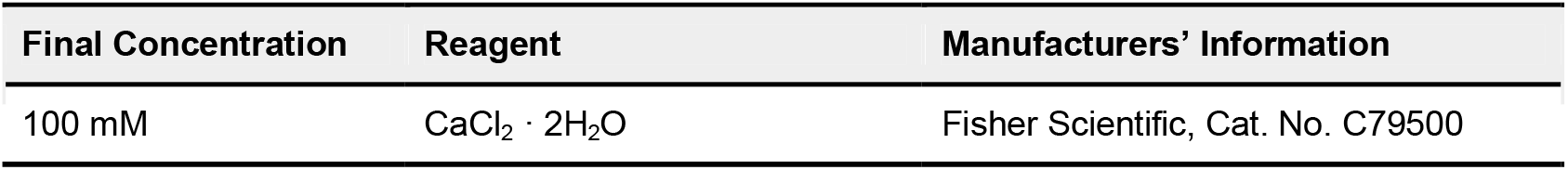
100 mM CaCl_2_.

**Table 3.**
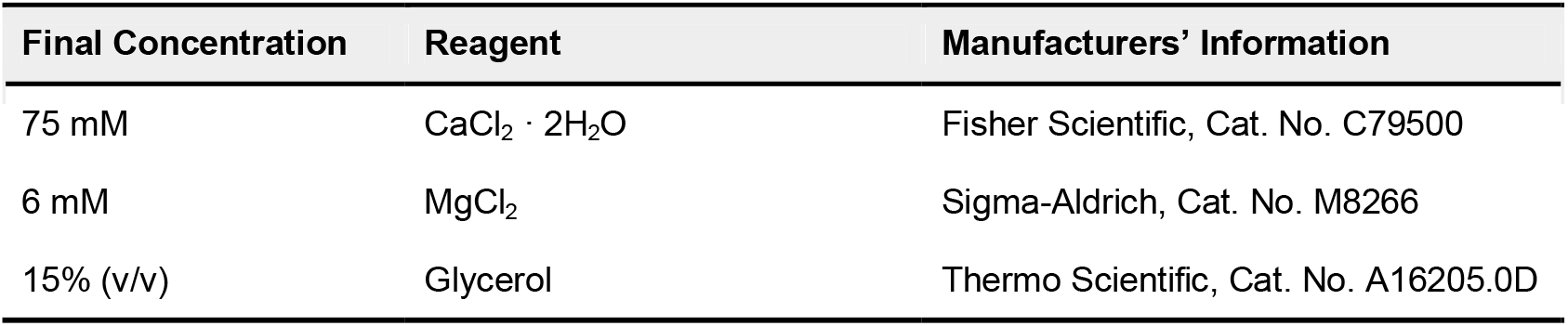
Transformation salts with Glycerol (TG salt solution) (7).

**Table 4.**
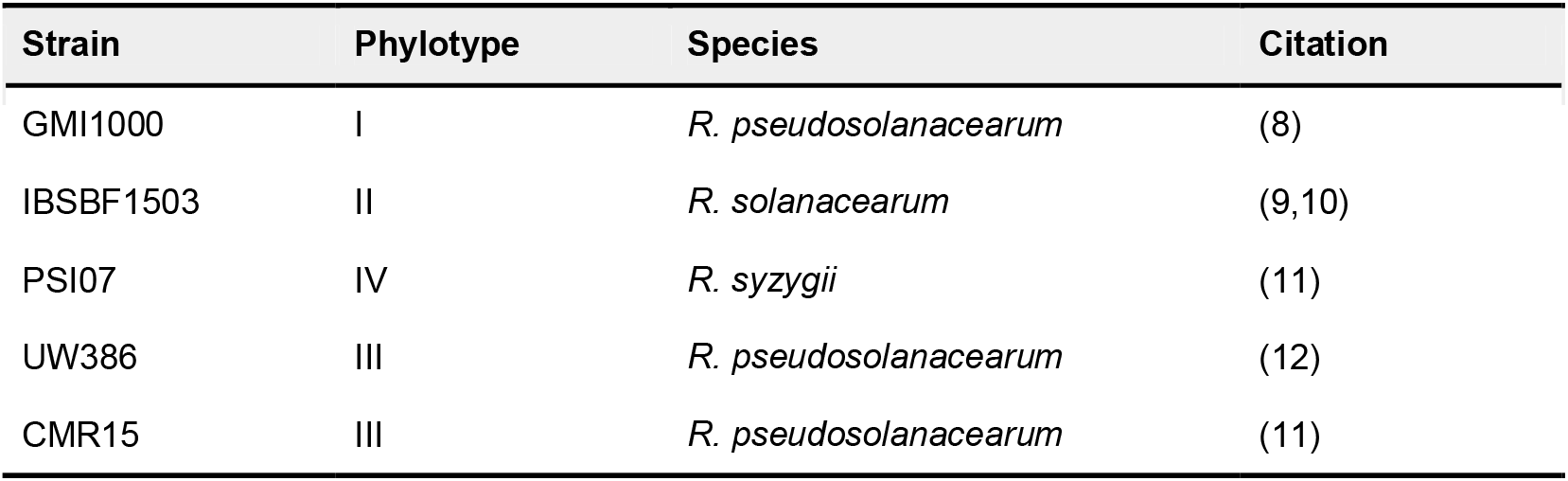
*Ralstonia* strains used in the development of the chemical competence protocol.

| Strain | Phylotype | Species | Citation |
| --- | --- | --- | --- |
| GMI1000 | I | *R. pseudosolanacearum* | (8) |
| IBSBF1503 | II | *R. solanacearum* | (9,10) |
| PSI07 | IV | *R. syzygii* | (11) |
| UW386 | III | *R. pseudosolanacearum* | (12) |
| CMR15 | III | *R. pseudosolanacearum* | (11) |

**Table 5.**
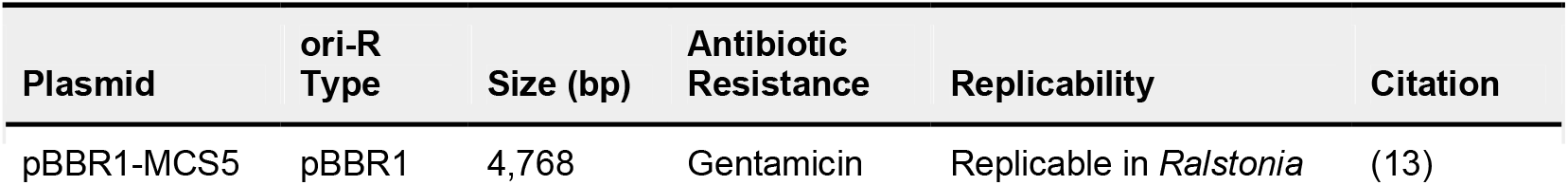
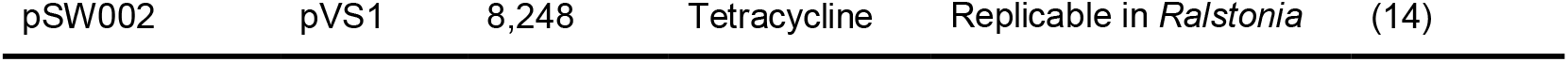
Plasmids used in the development of the chemical competence protocol.

For growing *E. coli*, we used LB Broth powder (Apex BioResearch Products, Cat. No. 11-121; 1.0% tryptone, 0.5% yeast extract, 1.0% sodium chloride), prepared as 25 g/L LB powder, with 15 g/L agar (Apex BioResearch Products, Cat. No. 20-248) for solid media, sterilized by autoclaving. Liquid cultures were grown in a shaking incubator at 37°C and 230 rpm. Cultures on agar plates were grown at 37°C. Concentrations and catalogs for antibiotics are listed in “Media”.

For cell density measurements in liquid, we used an Ultrospec 10 Cell Density Meter (Amersham Biosciences) to measure the absorbance at 600 nm (A_600_), based on the approximate conversion of an A_600_ value of 1 corresponding to a cell density of 5 · 10^8^ cfu/mL for *Ralstonia*.

### Chemically Competent *Ralstonia*

We used the described protocol to transform the *Ralstonia* strains GMI1000, IBSBF1503, PSI07, UW386, and CMR15 with the plasmids pBBR1-MCS5 and pSW002. For each strain, we performed 5 to 19 transformations using the final version of the protocol and calculated transformation efficiencies (**Figure 1**). For GMI1000 only, we also performed 12 to 28 transformations using the preliminary protocol, with both freshly prepared cells and cell suspensions stored at −80°C for four weeks (**Figure 1**). We used plasmid amounts ranging from 45.3 to 745.5 ng in volumes ranging from 0.8 to 15.0 μL.

**Figure 1.**
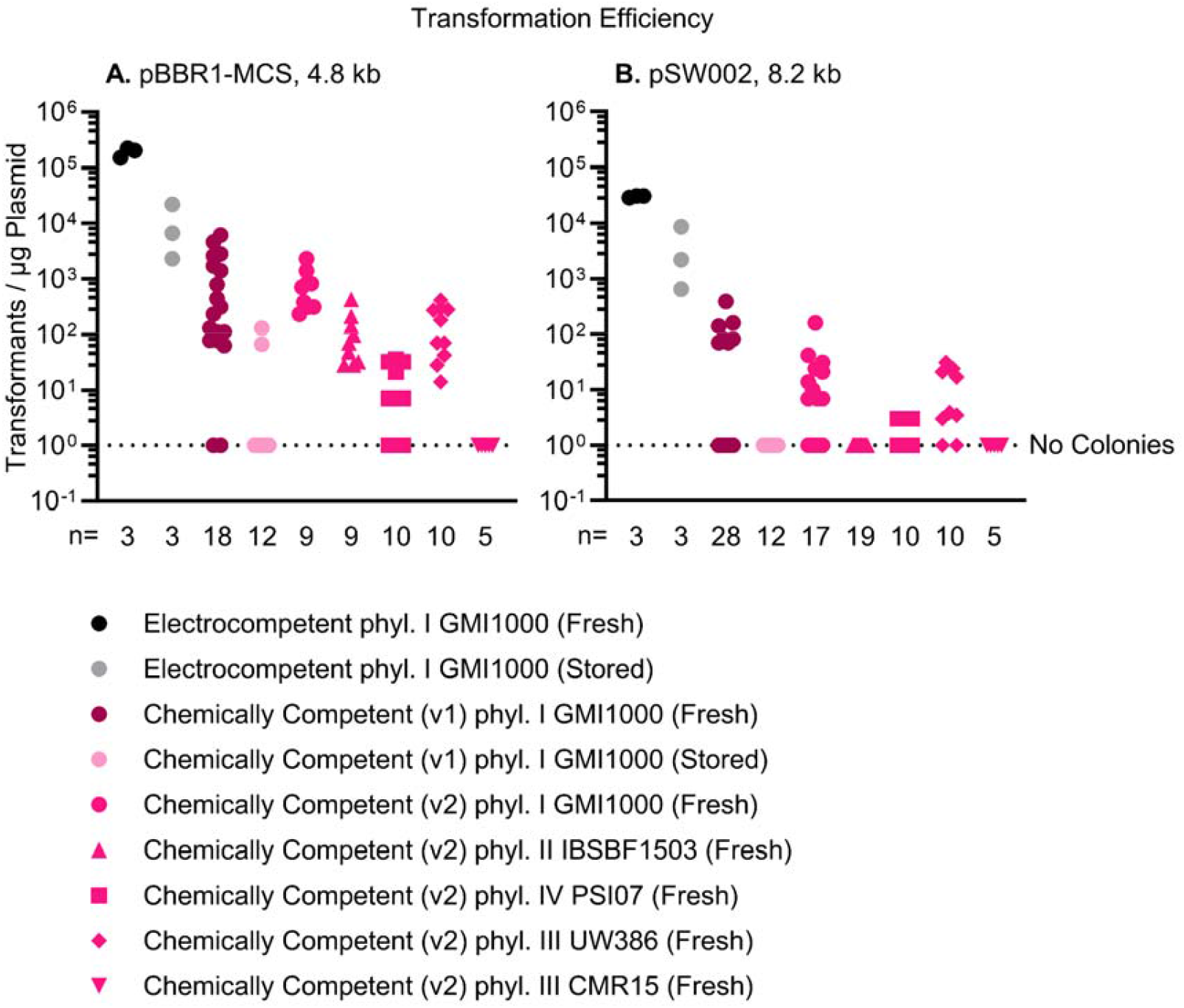
Transformation efficiencies for electroporation and chemical competence in selected *Ralstonia* strains (GMI1000, IBSBF1503, PSI07, UW386, and CMR15) using plasmids (**A**) pBBR1-MCS5 and (**B**) pSW002. The symbols represent individual transformations with symbol shape representing strain and symbol color representing the protocol variant. The dashed line indicates the limit of detection.

### Electrocompetent *Ralstonia*

To compare our chemical competence protocol to an existing transformation method in *Ralstonia*, we determined the transformation efficiency of electroporation of GMI1000 with the plasmids pBBR1-MCS5 and pSW002. We performed three replicates for each plasmid in two conditions: fresh cell suspensions and cell suspensions stored at −80°C for four weeks.

#### Culture Growth

GMI1000 was struck out from a −80°C glycerol stock onto CPG+TZC agar and incubated at 28°C for two days. A single colony was inoculated into 5 mL of CPG broth and incubated overnight in a shaking incubator at 28°C and 250 rpm. The A_600_ of the overnight culture was measured and diluted to an A_600_ of 0.001. Then, two replicate subcultures were prepared, each with 50 mL CPG broth and 1 mL of the A_600_ 0.001 suspension, and incubated in a shaking incubator at 28°C and 250 rpm for ∼21 hours until the subcultures reached an A_600_ of 0.48 and 0.52, respectively.

#### Cell Treatment

The subcultures were placed on wet ice for 10 min, then poured into two chilled 50 mL conical tubes. The subcultures were centrifuged at 5,000 x*g* for 10 min at 4°C, and the supernatant was poured off. The first pellet was resuspended in 10 mL of cold (4°C) 10% (v/v) glycerol by pipetting up and down. This glycerol-cell suspension was used to resuspend the second pellet, in order to pool the cells into one tube. The cells were pelleted by centrifugation and the supernatant was poured off. The cell pellet was resuspended in 6 mL of cold 10% glycerol by pipetting to mix. The cells were centrifuged and the supernatant was removed by pouring carefully to avoid disturbing the loose pellet. The cell pellet was resuspended in the residual glycerol by pipetting. Cells were aliquoted in 80 µL portions in chilled 1.5 mL microcentrifuge tubes on wet ice. At this point, the transformation steps were carried out for half of the aliquots to acquire the data for fresh cells. The remaining aliquots were moved to dry ice for 10 min before being moved to a −80°C freezer, where they were stored for four weeks before performing the transformation steps. These were used to acquire the data for stored cells.

#### Transformation

34.6 ng of pBBR1-MCS5 or 46.2 ng of pSW002 was added to an 80 µL aliquot of electrocompetent GMI1000 and mixed gently by pipetting, on ice. The mixture was transferred to a 1 mm electroporation cuvette and shocked in an Electroporator 2510 (Eppendorf) set to 900 V. Researchers in the Lowe-Power lab have had success varying electroporation from 900 to 1800 V, with anecdotes that higher voltages may increase electrotransformation efficiency. Immediately after the shock, 1 mL of CPG broth was added to the electroporation cuvette and mixed gently by pipetting up and down three times. The 1 mL mixture was transferred out of the electroporation cuvette and into a 2 mL microcentrifuge tube for outgrowth. This tube was shaken horizontally in a shaking incubator at 28°C and 250 rpm for 4 hours. Volumes (100 µL) of undiluted, 1:10 diluted, 1:100 diluted, and 1:1000 diluted outgrowth culture were plated onto pre-warmed CPG+TZC+antibiotic plates, spread using sterile glass beads. The plates were incubated for 3 days at 28°C. Then, colonies were counted and used to calculate transformation efficiency.

### Calculations

Here, transformation efficiency is defined as the number of transformants per μg of plasmid DNA. We calculated transformation efficiency (TE) using the following equation:

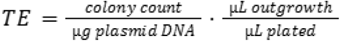

Results were graphed using GraphPad Prism v10.5.0 for Windows.

## Results

Freshly prepared electrocompetent GMI1000 had a median transformation efficiency of 200,000 (range: 150,000-220,000) transformants per μg of plasmid DNA with pBBR1-MCS5, and 31,000 (range: 29,000-31,000) transformants per μg of plasmid DNA with pSW002 (**Figure 1**). Stored electrocompetent GMI1000 had a median transformation efficiency of 6,600 (range: 2,300-22,000) transformants per μg of plasmid DNA with pBBR1-MCS5, and 2,200 (range: 650-8,700) transformants per μg of plasmid DNA with pSW002 (**Figure 1**).

Chemically competent GMI1000 that was freshly prepared using the preliminary protocol (Supplemental File S1) had a median transformation efficiency of 270 (range: 0-6,100) transformants per μg of plasmid DNA with pBBR1-MCS5, and 0 (range: 0-390) transformants per μg of plasmid DNA with pSW002 (**Figure 1**). Chemically competent GMI1000 that was stored after preparation using the preliminary protocol had a median transformation efficiency of 0 (range: 0-130) transformants per μg of plasmid DNA with pBBR1-MCS5 (**Figure 1**). There were no successful transformations with pSW002.

Chemically competent GMI1000 that was freshly prepared using the final protocol had a median transformation efficiency of 700 (range: 230-2,300) transformants per μg of plasmid DNA with pBBR1-MCS5, and 6.9 (range: 0-160) transformants per μg of plasmid DNA with pSW002 (**Figure 1**). Chemically competent IBSBF1503 that was freshly prepared using the final protocol had a median transformation efficiency of 70 (range: 28-420) transformants per μg of plasmid DNA with pBBR1-MCS5 (**Figure 1**). There were no successful IBSBF1503 transformations with pSW002. Chemically competent PSI07 that was freshly prepared using the final protocol had a median transformation efficiency of 7 (range: 0-35) transformants per μg of plasmid DNA with pBBR1-MCS5, and 0 (range: 0-3) transformants per μg of plasmid DNA with pSW002 (**Figure 1**). Chemically competent UW386 that was freshly prepared using the final protocol had a median transformation efficiency of 125 (range: 14-410) transformants per μg of plasmid DNA with pBBR1-MCS5, and 10.5 (range: 0-31) transformants per μg of plasmid DNA with pSW002 (**Figure 1**). For chemically competent CMR15 that was freshly prepared using the final protocol, there were no successful transformations with pBBR1-MCS5 or pSW002 (**Figure 1**).

Additionally, we tested this protocol and the preliminary protocol in GMI1000 with ∼10 kb, pUFR80-based plasmids (16) that are not replicable in *Ralstonia*, aiming to perform markerless gene knockouts through homologous recombination (Supplemental Figure S1). We used plasmid amounts of 1 to 6.3 μg. Successful transformations had efficiencies of 0.6 to 96.0 transformants per μg of plasmid DNA.

### Logic Underlying Protocol Development

The *Ralstonia* chemical competence protocol is adapted from a chemical competence protocol for *Pseudomonas aeruginosa* (7). The *P. aeruginosa* protocol treats cells with divalent cations by washing cells from overnight cultures in a magnesium chloride solution and then incubating, washing, and resuspending cells in the TG salt solution. The prepared cells are transformed through a heat shock at 37°C, followed by an outgrowth recovery and plating on selective media.

Due to a combination of literature review and iterative protocol testing, we adjusted the concentrations of calcium chloride and magnesium chloride in the wash solutions, incubation times of wash steps, and the temperature and number of heat shocks (17). For simplicity, testing was limited to methods that used calcium chloride and/or magnesium chloride as the inducer of chemical competence, but there are other chemicals (18–20) that have been shown to induce competence in other bacteria. Future protocol development could test whether alternative treatments like rubidium chloride (20), sepiolite (20), polyethylene glycol (18), dimethyl sulfoxide (18), polyamines (25), and combinations thereof are more efficient.

When we tested the use of calcium chloride and magnesium chloride in the first wash step, we found that calcium chloride alone was more effective than either magnesium chloride alone or a 1:1 combination of calcium chloride and magnesium chloride. This is consistent with existing data regarding chemical competence in other bacteria, which indicates that calcium ions tend to be the most effective cation for chemical competence (17).

It has been previously determined that extended incubation, up to 24 hours, of cells in calcium chloride improves transformation efficiency in *E. coli* (21). Since both wash solutions in this protocol contain calcium chloride, we decided to incubate the cells during each wash step. To keep the protocol within a reasonable time frame, the incubation times were limited to 20 and 15 min, respectively. However, longer incubation times, such as an overnight incubation, have the potential to further improve transformation efficiency.

Chemical competence protocols generally require one heat shock step; however, it has been shown that multiple heat shock/ice incubation cycles can result in increased transformation efficiency (22). We tested one, two, and three heat shock cycles for the chemical competence of *Ralstonia*. We also tested heat shock temperatures of 37°C, 42°C, 45°C, and 48°C. Our results indicated that three heat shocks at 42°C, 45°C, and 48°C all had similar transformation efficiencies; and one, two, and three heat shocks at 48°C all had similar transformation efficiencies (Supplemental Figure S2).

These adjustments collectively resulted in the final version of our protocol, but we also tested other factors, which are shown in our preliminary protocol (Supplemental File S1). The preliminary protocol used magnesium-enriched growth media instead of CPG, cells subcultured to exponential phase instead of overnight cultures, and took five days instead of four. The differences in transformation efficiency between the preliminary and final protocols were small enough that we opted to present the simpler and easier protocol above.

Previous studies have shown that the presence of magnesium ions in the media during overnight growth increases the transformation efficiency of chemically competent *E. coli* (23). Super Optimal Broth (SOB) and Super Optimal broth with Catabolite repression (SOC) both contain magnesium ions and are commonly used in the preparation and outgrowth of chemically competent *E. coli*, respectively (23). We adapted the SOB and SOC recipes to develop *Ralstonia*-specific versions, which we named ROB and ROC. The main modification was replacing the sodium chloride in SOB and SOC with the molar equivalent of potassium chloride in ROB and ROC because *Ralstonia* is sensitive to sodium chloride (24).

Additionally, SOB and SOC have the same basic components of LB (Lysogeny Broth) media, which is used to grow *E. coli*. Since *Ralstonia* is grown on CPG, we replaced the LB components with the corresponding CPG components in the ROB and ROC recipes.

However, we determined that transformation efficiency with ROB/ROC media was similar to standard CPG media. Thus, the final version of the protocol does not use ROB/ROC, but we included these recipes because some of the data in Figure 1 use cells grown in ROB/ROC (“Chemically Competent (v1)”; Supplemental File S1).

The physiological state of cells is known to influence transformation efficiency, so we tested *Ralstonia* from both mid-exponential phase subcultures and stationary phase overnight cultures. In our electrocompetence protocol for *Ralstonia*, we use cells that were subcultured and grown to mid-exponential phase. Similarly, when preparing chemically competent *E. coli* it is common to use subcultured cells grown to mid-exponential phase. Based on this, our preliminary chemical competence protocol used mid-exponential phase *Ralstonia* cells.

However, our testing showed that the subculturing step did not substantially impact transformation efficiency, so it was excluded from the final protocol.

## Author statements

### Author contributions

TCC: Analysis, Experiments / Investigation, Methodology, Validation, Visualization, Writing – original draft, NRG: Experiments / Investigation, Validation, Writing – review & editing, MLCA: Experiments / Investigation, Validation, Writing – review & editing, NNP: Experiments / Investigation, Validation, Writing – review & editing, TMLP: Conceptualization, Project administration, Resources, Supervision, Writing – review & editing.

## Acknowledgements

We would like to acknowledge the members of the Lowe-Power lab for valuable discussions.

