## Supplemental Figures for "A protocol for chemical competence in phytopathogenic *Ralstonia*"

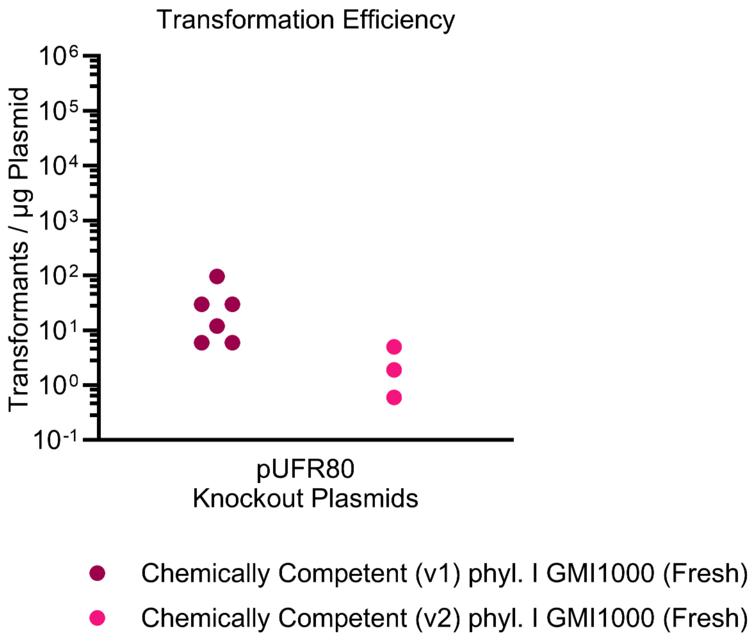


**Figure S1.** Transformation efficiencies for chemical competence in the *Ralstonia* strain GMI1000 using pUFR80 knockout plasmids. The symbols represent individual transformations with symbol color representing the protocol variant.

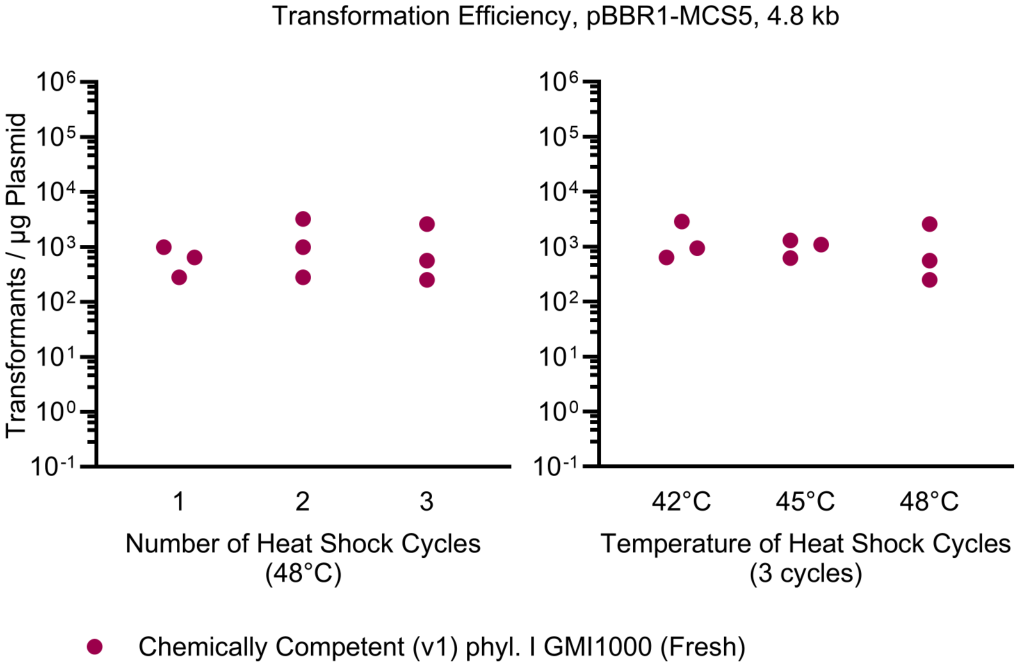


**Figure S2.** Transformation efficiencies for chemical competence in the *Ralstonia* strain GMI1000 using the plasmid pBBR1-MCS5 with varying heat shock cycles and temperatures. The symbols represent individual transformations. All transformations shown here were performed using the preliminary protocol.
