## Supplemental File S1 for "A protocol for chemical competence in phytopathogenic *Ralstonia*"

### Chemically Competent *Ralstonia*, Preliminary Protocol

#### Reagents

##### Media

**Table 1.** CPG rich medium (Kelman, 1954).

| Final Concentration | Reagent | Manufacturers' Information |
|---|---|---|
| 1 g/L | Casamino acids | Research Products International, Cat. No. C45000-500.0 |
| 10 g/L | Peptone | Apex BioResearch Products, Cat. No. 20-260 |
| 5 g/L | Glucose | Sigma-Aldrich, Cat. No. G8270 |
| 1 g/L | Yeast extract | Apex BioResearch Products, Cat. No. 20-254 |
| For solid media (CPG+TZC agar), add: | | |
| 15 g/L | Agar | Apex BioResearch Products, Cat. No. 20-248 |
| 2 mL/L | 1% (v/v) 2,3,5-triphenyl-2H-tetrazolium chloride (TZC) in water | Thermo Scientific, Cat. No. A10870.09 |

Media are sterilized by autoclaving. TZC and antibiotics should be added to molten CPG agar cooled to ~57°C.

**Table 2.** Super Optimal Broth (SOB) Modified for *Ralstonia* (ROB).

| Final Concentration | Reagent | Manufacturers' Information |
|---|---|---|
| 1 g/L | Casamino acids | Research Products International, Cat. No. C45000-500.0 |
| 15 g/L | Peptone | Apex BioResearch Products, Cat. No. 20-260 |
| 1 g/L | Yeast extract | Apex BioResearch Products, Cat. No. 20-254 |
| 12.5 mM | KCl | EMD, Cat. No. PX1405-1 |
| 10.0 mM | $MgCl_2$ | Sigma-Aldrich, Cat. No. M8266 |
| 10.0 mM | $MgSO_4 \cdot 7H_2O$ | Fisher Scientific, Cat. No. M80-500 |
| For Super Optimal broth with Catabolite repression (SOC) Modified for *Ralstonia* (ROC), add: | | |
| 20 mM (or 0.4% (w/v)) | Glucose | Sigma-Aldrich, Cat. No. G8270 |

ROB and ROC should be filter-sterilized using filters with a 0.22 μm pore size.

#### Solutions

**Table 3.** 100 mM $CaCl_2$.

| Final Concentration | Reagent | Manufacturers' Information |
|---|---|---|
| 100 mM | $CaCl_2 \cdot 2H_2O$ | Fisher Scientific, Cat. No. C79500 |

**Table 4.** Transformation salts with Glycerol (TG salt solution) (Chuanchuen *et al*., 2002).

| Final Concentration | Reagent | Manufacturers' Information |
|---|---|---|
| 75 mM | $CaCl_2 \cdot 2H_2O$ | Fisher Scientific, Cat. No. C79500 |
| 6 mM | $MgCl_2$ | Sigma-Aldrich, Cat. No. M8266 |
| 15% (v/v) | Glycerol | Thermo Scientific, Cat. No. A16205.0D |

Solutions should be filter-sterilized using filters with a 0.22 µm pore size. Solutions should be pre-chilled to 4°C prior to use.

### Procedure

After the written protocol, there are notes and recommendations for steps 5, 6, 10, 14, 17, 19, 24, and 25.

#### Culture Growth

1. Streak out *Ralstonia* strain on CPG+TZC agar and incubate at 28°C for 2 days or room temperature (20-22°C) for 3-4 days.
2. Inoculate a single colony into 5 mL CPG broth and incubate overnight in a shaking incubator at 250 rpm and 28°C.
3. Measure the absorbance at 600 nm ($A_{600}$) of overnight culture.
4. Dilute overnight culture to $A_{600}$ 1, then serial dilute to $A_{600}$ 0.001.
5. Add 100 µL of $A_{600}$ 0.001 culture to 5 mL ROB.
6. Incubate in a shaker incubator at 250 rpm and 28°C until the subculture reaches exponential phase ($A_{600}$ 0.2-0.6).

#### Cell Treatment

7. Perform all steps on ice except for centrifugations. Ensure that materials are pre-chilled: solutions and microcentrifuge tubes (1.5 mL and 2.0 mL). Set a water bath to 45°C.
8. Incubate subculture on ice for 10 min (make sure the culture tube is slightly buried in ice so that the full culture volume is chilled).
9. Add 1 mL overnight culture to a chilled 1.5 mL microcentrifuge tube.
10. Centrifuge for 30 s at 13,000 x*g* and pipette off the supernatant.
11. Repeat steps 9 and 10 in the same microcentrifuge tube to increase the cell mass in the pellet.
12. Resuspend cell pellet in 1 mL cold 100 mM $CaCl_2$ by pipetting to mix.
13. Incubate on ice for 20 min.
14. Centrifuge for 30 s at 13,000 x*g* and pipette off the supernatant.

15. Resuspend cell pellet in 1 mL cold TG salt solution by pipetting to mix.
16. Incubate on ice for 15 min.
17. Centrifuge for 30 s at 13,000 x*g* and pipette off the supernatant.
18. Resuspend cell pellet in 100 µL cold TG salt solution by pipetting to mix and keep the chemically competent cell suspension on ice until use.

**Transformation**

19. Add plasmid DNA and 100 µL of the chemically competent cell suspension to a chilled 2.0 mL microcentrifuge tube.
20. Incubate on ice for 15 min.
21. Heat shock in a water bath at 45°C for 2 min.
22. Incubate on ice for 15 min.
23. Repeat steps 21-22 twice for a total of three heat shocks and four ice incubations.
24. For the outgrowth, add 500 µL CPG broth and shake the tube horizontally for 4 hours in a shaking incubator at 250 rpm and 28°C.
25. Plate 100 µL of the outgrowth culture onto prewarmed plates containing CPG+TZC media with selective antibiotics. Incubate at 28°C for 2-5 days.

**Notes and Recommendations**

- Step 5: For larger culture volumes, the $A_{600}$ 0.001 culture volume and ROB volume can be proportionally scaled up.
- Step 6: For GMI1000, it takes ~20-21 hours for the subculture to be at an $A_{600}$ of 0.2-0.6. For slow growing strains, either plan to let them grow for more time or inoculate with $A_{600}$ 0.01 culture instead.
- Steps 10, 14, 17: Some cells will be lost after each centrifugation and supernatant removal, but this is not a problem when working with high density overnight cultures ($A_{600}$ > 2). If cells are centrifuged for longer than 30 s, we recommend using a temperature controlled centrifuge set to 4°C to prevent the cells from warming during centrifugation. We did not yield any transformants after six transformations from a single attempt to scale up "Cell Treatment" (Steps 3-13) by centrifuging in 50 mL conical tubes at 5,000 xg for 10 min at 4°C.
- Step 19: When using plasmids that are 8 kb or larger, we recommend using large amounts of plasmid DNA (≥ 1 µg) for higher rates of successful transformation. We chose to move cells to 2.0 mL microcentrifuge tubes in Step 19 because these tubes allow for better circulation of the culture during the outgrowth.
- Step 24: The outgrowth can be carried out for as little as 2 hours. Prolonged outgrowths in excess of 5 hours should be avoided so that each colony on the selection plate is derived from an independent genetic event.
- Step 25: The outgrowth culture can be diluted before plating, and a 1:10 dilution is usually sufficient. The outgrowth culture can also be concentrated by centrifuging and resuspending the cells in a smaller volume prior to plating.
